# Acetylcholine modulates cerebellar granule cell spiking by regulating the balance of synaptic excitation and inhibition

**DOI:** 10.1101/760223

**Authors:** Taylor R. Fore, Nathan Taylor, Nicolas Brunel, Court Hull

**Author notes:** Correspondence: Dr. Court Hull. Conflict of Interest Statement: The authors declare no competing financial interests.

## Abstract

Sensorimotor integration in the cerebellum is essential for refining motor output, and the first stage of this processing occurs in the granule cell layer. Recent evidence suggests that granule cell layer synaptic integration can be contextually modified, though the circuit mechanisms that could mediate such modulation remain largely unknown. Here we investigate the role of Acetylcholine (ACh) in regulating granule cell layer synaptic integration. We find that Golgi cells, interneurons that provide the sole source of inhibition to the granule cell layer, express both nicotinic and muscarinic cholinergic receptors. While acute ACh application can modestly depolarize some Golgi cells, the net effect of longer, optogenetically induced ACh release is to strongly hyperpolarize Golgi cells. Golgi cell hyperpolarization by ACh leads to a significant reduction in both tonic and evoked granule cell synaptic inhibition. ACh also reduces glutamate release from mossy fibers by acting on presynaptic muscarinic receptors. Surprisingly, despite these consistent effects on Golgi cells and mossy fibers, ACh can either increase or decrease the spike probability of granule cells as measured by non-invasive cell attached recordings. By constructing an integrate and fire model of granule cell layer population activity, we find that the direction of spike rate modulation can be accounted for predominately by the initial balance of excitation and inhibition onto individual granule cells. Together, these experiments demonstrate that ACh can modulate population-level granule cell responses by altering the ratios of excitation and inhibition at the first stage of cerebellar processing.

**Significance Statement:** The cerebellum plays a key role in motor control and motor learning. While it is known that behavioral context can modify motor learning, the circuit basis of such modulation has remained unclear. Here we find that a key neuromodulator, Acetylcholine (ACh), can alter the balance of excitation and inhibition at the first stage of cerebellar processing. These results suggest that ACh could play a key role in altering cerebellar learning by modifying how sensorimotor input is represented at the input layer of the cerebellum.

## Introduction

To coordinate movements and learn sensorimotor associations, the cerebellum must integrate signals across a variety of modalities and timescales (Giovannucci et al., 2017; Ito, 2006, 2012; Mauk and Buonomano, 2004; Wagner et al., 2017). These signals are conveyed to the cerebellum via mossy fiber projections that form excitatory connections onto a much larger network of granule cells. The divergence from mossy fibers to granule cells has long been hypothesized to enable expansion recoding that enhances pattern separation and increases the encoding capacity of the network (Albus, 1971; Cayco-Gajic et al., 2017; Gilmer and Person, 2017; Marr, 1969).

While classical models of granule cell layer function posit rigid integration rules dictated by the anatomical pattern of mossy fiber to granule cell connectivity, recent studies have demonstrated that the granule cell layer can be contextually modified during behavior. Specifically, granule cell responses to certain sensory modalities can be suppressed during locomotion (Ozden et al., 2012). Moreover, associative learning can be enhanced during locomotion via circuit modulation thought to occur in the granule cell layer (Albergaria et al., 2018). Such contextual modification of granule cell responses may allow for both the enhancement of behaviorally relevant stimuli that should be learned, as well as suppression of information that should not be learned. However, the cellular mechanisms that allow such context-dependent regulation are largely unknown.

Throughout the brain, neuromodulators play a key role in contextual circuit regulation (Froemke et al., 2013; Kuchibhotla et al., 2017; Letzkus et al., 2011). In the cerebellum, there are prominent ACh projections that terminate in the granule cell layer (Jaarsma et al., 1997). Moreover, immunohistochemical studies have shown both nicotinic and muscarinic receptors in the cerebellar cortex (Dominguez de Toro et al., 1997; Dominguez del Toro et al., 1994; Nakayama et al., 1997; Neustadt et al., 1988; Turner and Kellar, 2005). These anatomical observations suggest that ACh could be a key regulator of cerebellar processing and cerebellar-dependent behaviors. Indeed, behavioral studies have demonstrated cholinergic enhancement of the OKR and VOR reflexes (Prestori et al., 2013; Tan and Collewijn, 1991, 1992).

To test how acetylcholine acts to modulate granule cell layer processing and synaptic integration, we have investigated both cell autonomous and circuit level effects of ACh by recording from granule cell layer neurons in an acute, in-vitro brain slice preparation. We find that ACh predominantly leads to a prolonged suppression of Golgi cell activity via muscarinic receptor activation, in turn reducing both tonic and evoked synaptic inhibition onto granule cells. In addition, activation of presynaptic muscarinic receptors on mossy fibers leads to a reduction in granule cell excitation. Together, the coincident reduction in excitation and inhibition increases spike probability in some granule cells, while reducing spike probability in others. A population level integrate and fire model of granule cell layer synaptic processing reveals that the direction of modulation depends on the relative balance of excitation and inhibition of individual granule cells. Specifically, we find that the activity of granule cells with the most inhibition is preferentially enhanced by ACh, whereas the activity of granule cells with little inhibition is largely suppressed. Thus, these data suggest that ACh can act to enhance the reliability of granule cells that are significantly inhibited in response to specific mossy fiber input. Such modulation would be well-suited to enhance the responses of granule cells that receive stimulus specific inhibition without expanding the overall population response.

## Materials and Methods

### Acute Slices and Recordings

Acute sagittal slices (250 μm) were prepared from the cerebellar vermis of Sprague Dawley rats (20–25 days old, males, Charles River) and ChAT-IRES-Cre mice (B6;129S6-*Chat^tm2(cre)Lowl^*/J, Jackson Labs, P40-60, males and females). Slices were cut in an ice-cold potassium cutting solution (Dugue et al., 2005) consisting of (in mM): 130 K-gluconate, 15 KCl, 0.05 EGTA, 20 HEPES, 25 glucose (pH 7.4, 320 mmol/kg), and were transferred to an incubation chamber containing artificial CSF comprised of (in mM): 125 NaCl, 26 NaHCO_3_, 1.25 NaH_2_PO_4_, 2.5 KCl, 2 CaCl_2_, 1 MgCl_2_, and 25 glucose (pH 7.3, 320 mmol/kg). A NMDA antagonist, *R*-CPP (2.5 μM, Tocris), was added to the potassium cutting solution to enhance cell survival. Slices were incubated at 32° C for 20 minutes, and then kept at room temperature for up to 7 hours. All solutions were saturated with 95% O_2_ and 5% CO_2_. All procedures were performed according to guidelines approved by the Duke University Institutional Animal Care and Use Committee.

Visually guided (SliceScope Pro 2000 with Dodt-gradient contrast and water-immersion 60x objective, Scientifica) whole-cell recordings were obtained using a Multiclamp 700B (Axon Instruments) with thick-walled borosilicate glass patch pipettes (Granule cells, whole cell: 5-7 MΩ, Granule cells, cell attached: 10-14 MΩ, Golgi cells: 2-4 MΩ; 1.5 mm OD, 0.84 mm ID, World Precision Instruments). Electrophysiological recordings were digitized at 20 kHz (Digidata 1440A, Axon Instruments), filtered at 10 kHz, and performed at 32-33 °C. Glass monopolar electrodes (1 MΩ) filled with aCSF, in conjunction with a stimulus isolation unit (ISO-Flex, A.M.P.I.), were used for extracellular stimulation of the mossy fiber tract. For cell-attached recordings, only cells with an initial spike probability in control conditions between approximately 20% and 60% were recorded to allow for increases or decreases in spike rate during pharmacological manipulations.

Spontaneous and evoked IPSCs were recorded at the EPSC reversal potential (+10 mV). Evoked excitatory postsynaptic currents (eEPSC) were recorded at a holding potential of −70 mV and −60 mV for granule cells and Golgi cells, respectively. Voltage-clamp recordings of IPSCs and EPSCs were collected using a cesium-based internal solution containing (in mM): 140 Cs-gluconate, 15 HEPES, 0.5 EGTA, 2 TEA-Cl, 2 MgATP, 0.3 NaGTP, 10 Phosphocreatine-Tris_2_, 2 QX-314 Cl, pH was adjusted to 7.2 with CsOH, resulting in a final osmolality of 316 mmol/kg. Current-clamp and voltage-clamp recordings of Golgi cells were performed using a potassium-based internal solution containing (in mM): 150 K-gluconate, 3 KCl, 10 HEPES, 0.5 EGTA, 3 MgATP, 0.5 NaGTP, 5 Phosphocreatine-Tris_2_, 5 Phosphocreatine-Na_2_, pH was adjusted 7.2 with KOH, osmolality: 315 mmol/kg. Membrane potentials were not corrected for the liquid junction potential. Series resistance was monitored during voltage-clamp recordings with a 5 mV hyperpolarizing pulse, and only recordings that remained stable over the period of data collection were used. All drugs were purchased from Abcam or Tocris.

### Virus Injections and Optogenetic Experiments

Expression of the optogenetic actuator Chronos in ChAT+ cerebellar projection neurons was achieved using retro-orbital injection (2-10 μ|L)(Yardeni et al., 2011) of AAV-PHP.eb-Syn-FLEX-rc[Chronos-tdTomato] into ChAT-IRES-Cre mice (B6;129S6-*Chat^tm2(cre)Lowl^*/J, Jackson Labs) at P0-P2. Recordings with optogenetic stimulation were subsequently performed between P40-P60. Labeled axons in these experiments were observed throughout the cerebellar vermis, including both thin fibers and mossy fiber-like axons. Mossy fiber-like axons were most dense in lobules VIII-X. Experiments were performed in any vermis lobule where labeling was evident as visualized by Td-Tomato expression during experiments. Optogenetic stimulation was performed using full-field illumination (50 Hz, 2-3 sec, 10-20 mW) with a 450 nm laser (Optoengine, MGL-III-450).

### Data Analysis

IPSCs and EPSCs were analyzed using Clampfit, Igor Pro, and Mini Analysis software (v6.0.3, Synaptosoft Inc.) using a 3 kHz low pass Butterworth filter. Detection thresholds for single trial responses were set to 5x (IPSCs) or 2.5x (EPSCs) greater than the baseline RMS noise level. Synaptic potency was measured as the amplitude on all trials where EPSCs or IPSCs were successfully evoked, excluding failures. Mossy fiber paired-pulse ratios were determined using an inter-stimulus interval (IS) of 40 ms and measured from an average waveform across 30 consecutive trials. For voltage-clamp recordings of nicotinic and muscarinic currents, peak current amplitudes were measured from the subtracted average current for each cell. For cell attached recordings, spike probabilities were calculated in 1 ms bins, and normalized to the maximum spike probability on a per cell basis. Spike rates were calculated across all three stimuli in a 45 ms window and normalized to control. Data are reported as mean ± SEM (unless otherwise noted), and statistical analysis was carried out using custom R package (available at github.com/trfore/MAtools) and Clampfit (Molecular Devices). Data was tested for homoscedasticity using Brown-Forsythe test and for normality via quantile-quantile plots. For heteroscedastic data, we applied a repeated measures ANOVA with Dunnett’s post hoc test; additionally, sphericity was not assumed and a Greenhouse-Geisser correction was applied. Alternatively, a one-way ANOVA with Tukey post hoc was used. Significant differences are denoted as follows: *** = p < 0.001, ** = p < 0.01, and * = p < 0.05.

### Modelling

The granular layer model was simulated with the Brian simulator (http://briansimulator.org). The structure of the network was adapted from Solinas et al. (Solinas et al., 2010), which aims to recreate a functionally relevant cube of the cerebellar granular layer with 100 μm edge length. The model comprised 315 mossy fibers, 4096 granule cells, and 27 Golgi cells. Probabilistic synapses were formed using the convergence ratios in Table 1, with the probability of a particular presynaptic neuron making a connection with a particular postsynaptic neuron defined as p = (Conv. ratio) / (Total no. presynaptic neurons). There were no spatial constraints on synapse formation.

**Table 1.**
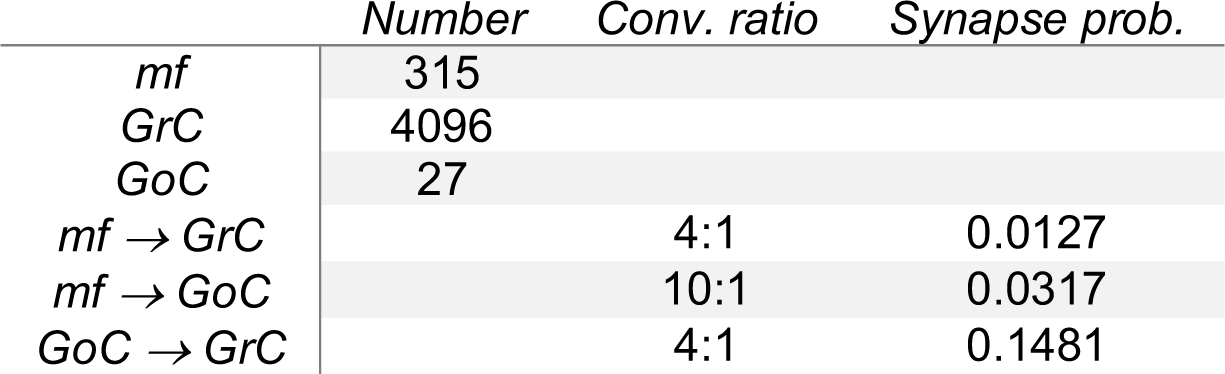

|  | *Number* | *Conv. ratio* | *Synapse prob.* |
| --- | --- | --- | --- |
| *mf* | 315 |  |  |
| *GrC* | 4096 |  |  |
| *GoC* | 27 |  |  |
| *mf* → *GrC* |  | 4:1 | 0.0127 |
| *mf* → *GoC* |  | 10:1 | 0.0317 |
| *GoC* → *GrC* |  | 4:1 | 0.1481 |

Granule and Golgi cells were modeled as conductance-based leaky integrate-and-fire neurons with sub-threshold membrane dynamics governed by:

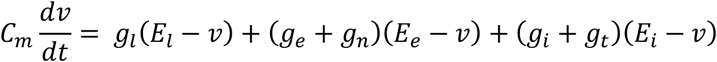

for granule cells, and

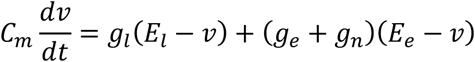

for Golgi cells, where *C_m_* is the membrane capacitance; *v* is the membrane potential; *E_l_*, *E_e_* and *E_i_* are leak, excitatory, and inhibitory reversal potentials; *g_l_*, *g_e_*, and *g_i_* are leak, excitatory, and (phasic) inhibitory conductances; *g_t_* is the (fixed) tonic inhibitory conductance in granule cells; and *g_n_* is a stochastically fluctuating excitatory conductance described by an Ornstein-Uhlenbeck process,

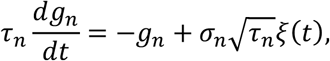

where *ξ*(*t*) is a white noise with unit variance density. Table 2 lists the fixed parameters for each cell type in control conditions. In control, Golgi cells fired spontaneously on average between 1 and 6 Hz. This spontaneous activity was achieved by setting the Golgi cell leak reversal +_)_ near spike threshold and allowing the stochastic excitatory current to occasionally drive the cell above threshold. Application of GABAzine was simulated by eliminating both tonic and phasic inhibition onto granule cells, without changing Golgi cell excitability. Tonic release of acetylcholine was simulated by suppressing Golgi cell activity (viz., by setting the leak reversal back to the resting membrane potential) and reducing the magnitude of mossy fiber excitation onto both granule and Golgi cells by 54.79% and 54.22% of their control values, respectively, as per experimental results. Tonic inhibition was reduced differentially across granule cells by a fraction drawn from a Gamma distribution with mean 0.6 and standard deviation 0.07.

**Table 2.**

| | $v_r$ | $v_{th}$ | $C_m$ | $g_l$ | $g_t$ | $E_l$ | $E_e$ | $E_i$ | $\sigma_n$ | $\tau_n$ | $\tau_e$ | $\tau_i$ |
| --- | --- | --- | --- | --- | --- | --- | --- | --- | --- | --- | --- | --- |
|  | (mV) | (mV) | (pF) | (nS) | (nS) | (mV) | (mV) | (mV) | (nS) | (ms) | (ms) | (ms) |
| GrC | -75 | -55 | 3.1 | 0.2 | 1 | -75 | 0 | -75 | 0.05 | 20 | 12 | 20 |
| GoC | -55 | -50 | 60 | 3 | 0 | -51 | 0 | -75 | 0.1 | 20 | 12 | - |

Three conditions (control, GABAzine, muscarine) were each simulated for 100 trials, each trial running for two seconds. Trials were simulated for one second without mossy fiber input to allow for decorrelation of GrC and GoC populations. Mossy fiber stimulation was modeled as three spikes delivered to a random selection of 32 mossy fibers at 1.01, 1.02, and 1.03 s. The same 32 mossy fibers were stimulated across all trials of an individual simulation. Each mossy fiber was assigned a mean synaptic weight from a gamma distribution with a mean and standard deviation of 0.1 nS. Following a presynaptic mossy fiber spike, excitatory postsynaptic conductance *g_e_* in both granule and Golgi cells was instantaneously increased by a synaptic weight drawn from a gamma distribution with mean and standard deviation equal to that mossy fiber’s assigned mean. A presynaptic Golgi cell spike elicited a fixed instantaneous jump in the inhibitory conductance *g_i_* on the postsynaptic granule cell with a 2 ms delay. Synaptic conductances decayed following the equations:

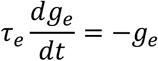

And

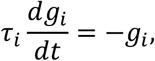

where *τ_e_* and τ_i_ are decay time constants specific to excitation and inhibition in each cell type. Upon reaching threshold *v_th_*, the membrane potential was immediately reset to its resting rate *v_r_*, and held there for an absolute refractory period of 2 and 10 ms for granule and Golgi cells, respectively.

Excitatory and inhibitory conductance traces were recorded for each granule cell during the simulation, as well as spike times. Spike probabilities were calculated in 1 ms bins over the 100 trials, and normalized to the peak spike probability on a per cell basis. Only granule cells with active mossy fiber input and a nonzero spike probability ratio in control were considered for subsequent analysis. To match with experimental criteria, we further selected for those cells whose peak spike probability to the second stimulus did not exceed 0.6, and whose peak spike probability to the third stimulus did not exceed that to the second. Cells categorized as ‘increasing’ spike probability in muscarine had a mean normalized spike probability greater than 1, while ‘decreasing’ cells had a mean normalized spike probability less than 1. Average conductance was calculated by integrating each conductance trace over a 60 ms period starting 10 ms before the first stimulus and ending 30 ms after the final stimulus, then dividing by that interval.

## Results

### Cholinergic modulation of Golgi Cells

Anatomical studies have reported nicotinic and muscarinic receptors in the cerebellar cortex (Jaarsma et al., 1997). Additionally, functional studies have found α7-nAChRs (Prestori et al., 2013), α4β2-nAChRs, and M3-mAChRs on granule cells; however, α4β2-nAChRs are reportedly developmentally down-regulated (Didier et al., 1995) and the M3-mAChRs are expressed in ∼15% of granule cells in lobules IX-X (Takayasu et al., 2003). To test for functional cholinergic receptors in the adult cerebellar granule cell layer, we performed whole-cell recordings from Golgi and granule cells in the presence of synaptic receptor antagonists (NBQX 5 μM, R-CPP 2.5 μM, and SR95531 5 μM), and focally applied acetylcholine (ACh 500 μM, 200 ms) via a second pipette positioned in close proximity (∼5-20 μm) to the recorded somata.

Golgi cells are the primary inhibitory interneuron in the granule cell layer, providing the main source of synaptic inhibition to granule cells. Because Golgi cells fire spontaneously between 1 to 20 Hz (Forti et al., 2006), we first tested for acetylcholine-mediated regulation of their spontaneous firing (**Figure 1A**). Brief application of ACh caused a transient increase in the firing rates of 7 of 10 recorded Golgi cells (5.3 ± 1.2 fold, n = 7 of 10, **Figure 1B, C, D**). However, in all recorded Golgi cells, there was a longer lasting suppression of firing (0.13 ± 0.06 fold, n = 10). To test what synaptic currents were responsible for these changes in spontaneous firing rate, we recorded in voltage-clamp configuration from a subset of the same cells. Focal application of ACh revealed a prolonged outward current that was blocked by the selective type II muscarinic receptor antagonist AF-DX 116 (1 μM)(16.1 ± 2.6 pA, n = 7, **Figure 1E,F**), as well as a transient inward current that was blocked by the non-selective nicotinic receptor antagonist MMA (10 μM) (40.3 ± 11.4 pA, n = 7, **Figure 1E,F**). In contrast, granule cell recordings revealed no change in membrane potential in response to ACh application in current-clamp, and no nicotinic nor muscarinic receptor-activated currents in voltage-clamp (n = 24, **Figure 1 G-J**).

**Figure 1.**
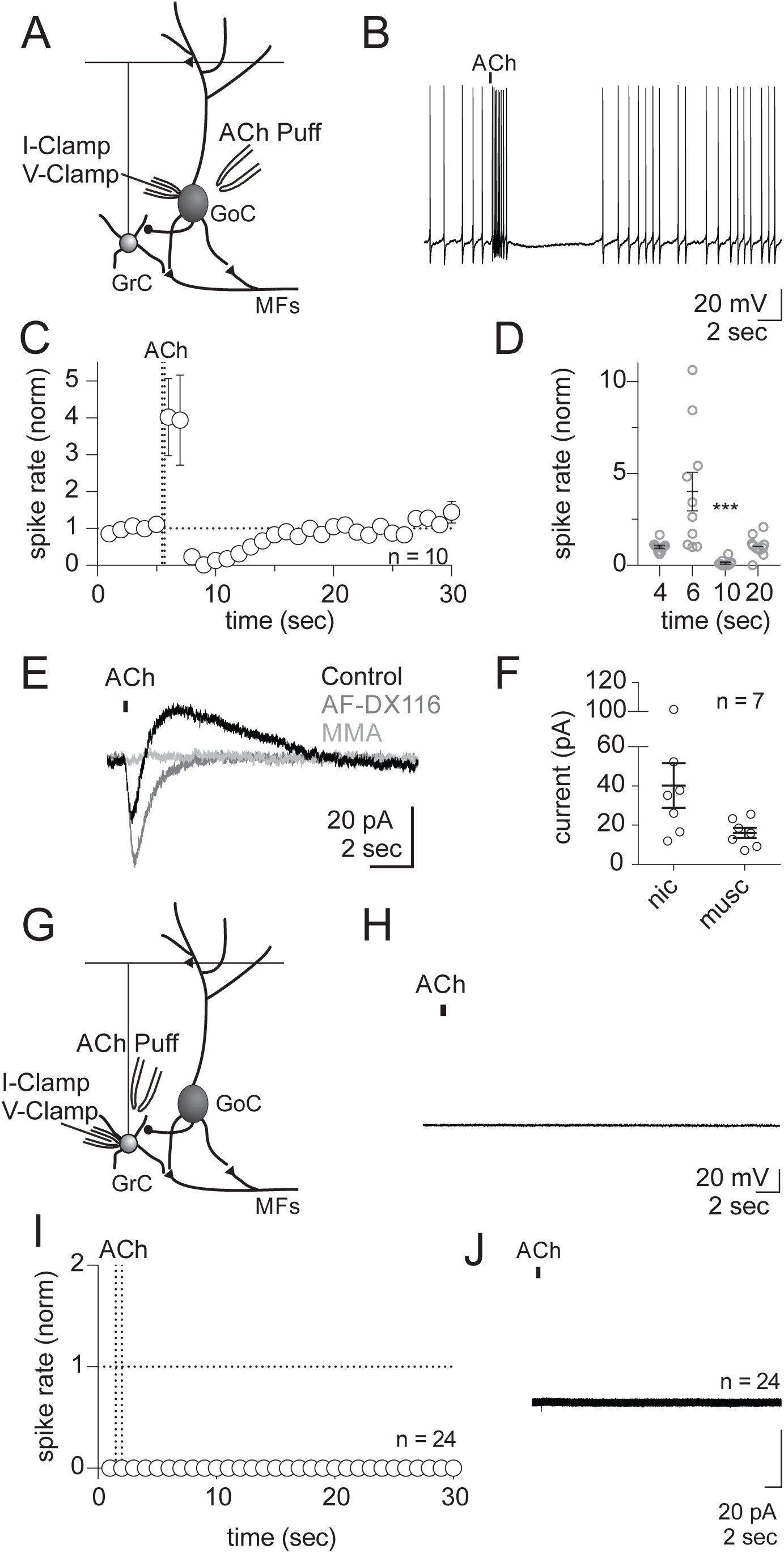
Acetylcholine activates nicotinic and muscarinic (M2) receptors in Golgi cells but does not act on granule cells. **A)** Schematic showing intracellular recording of Golgi cells during focal application of acetylcholine chloride (ACh). A glass pipette containing 500 μM ACh was placed in close proximity to the soma. **B)** Representative current-clamp recording of spontaneous activity in the presence of glutamatergic (NBQX 5 μM, CPP 2.5 μM) and GABAergic (SR-95531 5 μM) antagonist. A 200 ms pulse of ACh evokes a transient increase in spontaneous activity followed by a pause. **C-D)** Normalized spike rate across stimulus presentations (normalized to the 5 sec period prior to ACh stimulus, 1 sec bins, n = 10). **C)** Time course of ACh mediated change in spike rate (group average). **D)** Average spike rate by cell (grey), group average (black, mean ± SEM; ANOVA, F_1.052, 9.471_ = 9.679, P = 0.0111, n = 10) at 4 sec (0.99 ± 0.09 norm), 6 sec (4.0 ± 1.1 norm, P = 0.0517), 10 sec (0.1 ± 0.06 norm, P = 0.0004), and 20 sec (1.1 ± 0.2 norm, P = 0.9658). **E)** Voltage-clamp recordings from the same cell in B. A 200 ms pulse of ACh evokes an outward current that is blocked by a selective muscarinic M2 antagonist (AF-DX116 1 μM), and an inward current that is blocked by a non-selective nicotinic antagonist (MMA 10 μM). **F)** Peak amplitudes from voltage-clamp recordings at −60 mV. Amplitudes were determined using the subtracted average current for each cell (nicotinic: 40.3 ± 11.4 pA, muscarinic: 16.1 ± 2.6 pA, n = 7). **G)** Schematic showing intracellular recording of granule cells during focal application of acetylcholine chloride (ACh). A glass pipette containing 500 μM ACh was placed in close proximity to the soma. **H)** Representative current clamp trace from a granule cell in the presence of glutamatergic (NBQX 5 μM, CPP 2.5 μM) and GABAergic (SR-95531 5 μM) antagonist. A 500 ms pulse of ACh fails to evoke change in the membrane potential. **I)** Average spike rate across all cells (n = 24). **J)** Grouped voltage-clamp trace from all cells (n = 24).

Focal application of ACh onto Golgi cells suggests the possibility of long-term suppression during periods of ongoing acetylcholine release when nicotinic receptors are desensitized. To test this, we simulated tonic release of acetylcholine by bath application of muscarine (10 μM) in the presence of TTX (0.5 μM) to block spontaneous pacemaking (**Figure 2A**). In these experiments, muscarine produced a prolonged hyperpolarization that lasted the duration of the agonist application, and was abolished in the presence of the non-selective muscarinic receptor antagonist atropine (5 μM). To test whether such long-term modulation could be achieved by endogenous release of ACh, we expressed the optogenetic actuator chronos in cerebellar projecting cholinergic neurons (methods), and stimulated labeled axons in cerebellar vermis using 2-3 second pulse trains (50 Hz) of blue light (**Figure 2 B-E**). This stimulation protocol produced a modest depolarizing current during the stimulation, but predominantly resulted in a long-lasting, atropine sensitive outward current that emerged after the stimulus and persisted for 10s of seconds (**Figure 2D,E**). Overall, these finding suggest that ongoing bouts of granule cell layer ACh release can activate M2 muscarinic receptors on Golgi cells that produces a long-lasting, net suppression of these interneurons.

**Figure 2.**
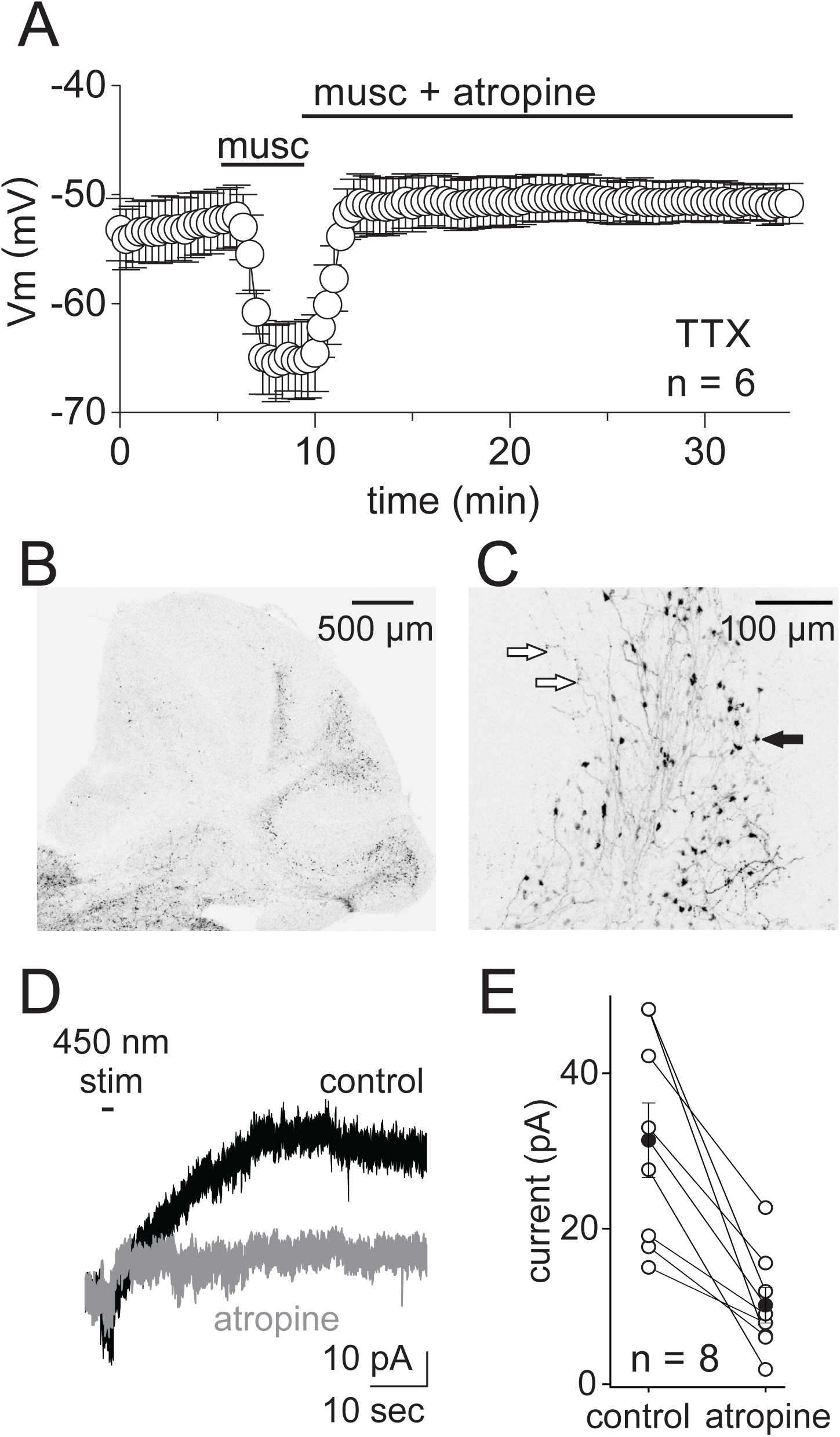
Activation of Muscarinic Receptors on Golgi Cells Hyperpolarizes the Membrane. **A)** Whole cell current-clamp recordings from Golgi cell in the in the presence of a Na^+^ channel antagonist, tetrodotoxin (TTX, 0.5 μM), to block spontaneous firing. Bath application of muscarine hyperpolarizes the membrane potential (musc: −65.2 ± 3.57 mV, n=6). **B-C)** Expression of chronos-TdTomato in ChAT+ cerebellar projections. **C)** Expanded view of the cerebellar cortex from B). Open arrows indicate thin, beaded fibers, closed arrow indicates mossy fiber-like projection. **D)** Average voltage-clamp recording from an example Golgi cell (Vm = −55 mV) in response to optogenetic stimulation (450 nm, 50 Hz, 15 mW, 2 sec) in control (black) and following atropine application (5 μM, gray) (control average = 31.4 ± 4.8 pA, atropine average = 10.2 ± 2.3 pA, n = 8, p = 0.002, paired t-test). **E)** Summary of Golgi cell optogenetic stimulation experiments.

### Cholinergic regulation of granule cell tonic inhibition

Golgi cells are the main source of granule cell synaptic inhibition (Eccles et al., 1966; Farrant and Brickley, 2003; Kanichay and Silver, 2008). Specifically, GABA released from Golgi cells activates two types of granule cell GABA_A_ receptors to regulate granule cell excitability (Brickley et al., 1996; Crowley et al., 2009; Duguid et al., 2015; Duguid et al., 2012). Based on our measurements of muscarinic receptor-mediated Golgi cell suppression, we next tested for changes in tonic granule cell synaptic inhibition during simulated bulk release of acetylcholine using bath application of muscarine (**Figure 3**). Bath application of muscarine resulted in a significant decrease in both spontaneous IPSC (sIPSC) frequency (83.7 ± 2.9% decrease, control: 4.5 ± 0.8 Hz, muscarine: 0.8 ± 0.3 Hz, n=11, P<0.0001, **Figure 3B**) and a tonic inhibitory current mediated by non-desensitizing GABA_A_ receptors (54.3 ± 3.3% decrease, control: 8.85 ± 1.08 pA, muscarine: 4.1 ± 0.7 pA, P<0.0001, **Figure 3C**). These effects were reversed by atropine (sIPSC frequency: 4.9 ± 0.9 Hz, holding current: 7.5 ± 1.0 pA, **Figure 3B-D**), suggesting that M2 receptor-mediated suppression of Golgi cell firing can significantly reduce tonic granule cell synaptic inhibition.

**Figure 3.**
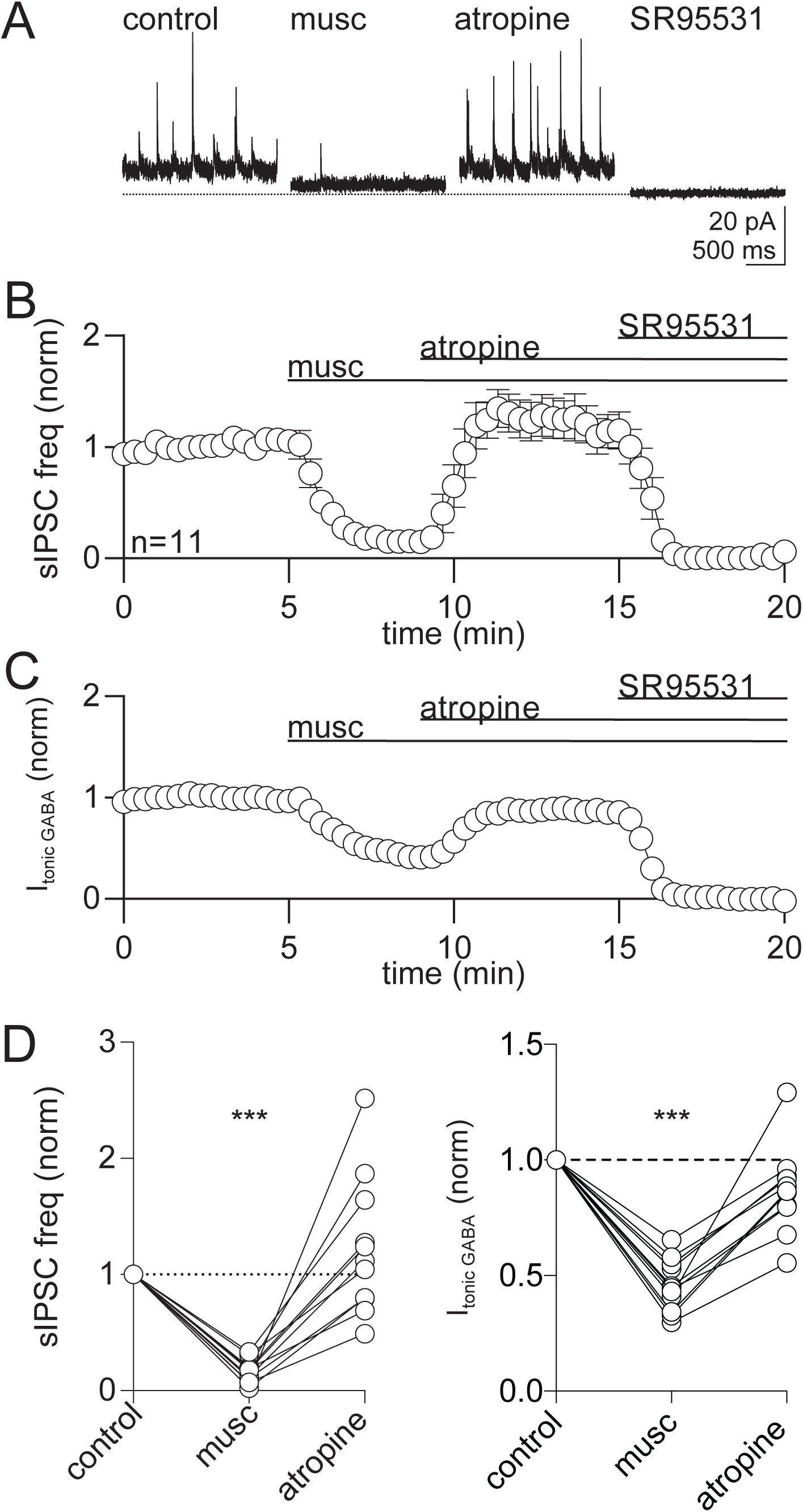
Activation of muscarinic M2 receptors on Golgi cell decreases phasic and tonic inhibition in granule cells. **A)** Example whole-cell voltage clamp recording from a granule cell held at the reversal potential of excitation (∼10 mV) to record GABA_A_ receptor-mediated phasic and tonic inhibition (measured using sIPSC frequency and holding current, respectively). Dashed horizontal line indicates no holding current. **B)** Grouped average spontaneous IPSC frequency across time in the presence of muscarine (musc, 10 μM), atropine (5 μM), and GABA_A_ antagonist (SR-95531 5 μM; mean ± SEM, n = 11). **C)** Grouped average holding current across time. **D)** *Left*, Summary of the change in sIPSC frequency for individual cells. Quantified using a 2-minute window (control: 2-4’, muscarine: 7-9’, atropine: 12-14’, values normalized to control; ANOVA, F_1.041, 10.41_=29.49, P = 0.0002, n = 11). Group average for each condition: muscarine (0.2 ± 0.0 norm, P<0.0001); atropine (1.2 ± 0.2 norm, P = 0.3773). *Right*, Summary of the change in holding current (ANOVA, F_1.565, 15.65_= 65.96, P<0.0001) for individual cells in muscarine (group average: 0.5 ± 0.0 norm, P<0.0001) and atropine (group average: 0.9 ± 0.1 norm, P = 0.0681). sIPSC and tonic inhibition was confirmed using the GABA_A_ antagonist (SR95531 5 μM).

### Activation of Golgi Cell M2 receptors induces spike rate plasticity

In a subgroup of granule cells, sIPSC frequency increased following bath application of muscarine (1.7 ± 0.2 of control, n=5 of 11, P = 0.015, **Figure 3D**). These data suggest an increase in spontaneous Golgi cell firing rates following muscarine application. Previous work has demonstrated that Golgi cell pacemaking can undergo a form of long-term firing rate potentiation following extended membrane hyperpolarization (Hull et al., 2013). To test whether activation of M2 receptors could provide an endogenous mechanism for inducing this plasticity, we performed whole-cell current clamp recordings from Golgi cells in the presence of synaptic receptor antagonists (NBQX 5 μM, CPP 2.5 μM, SR95531 5 μM) while simulating bulk acetylcholine release via bath application of muscarine (10 μM). Muscarine reliably suppressed GoC spiking throughout the application period (baseline: 4.2 ± 0.6 Hz; musc: 0.5 ± 0.3 Hz; **Figure 4A**). Following subsequent atropine application, Golgi cell firing resumed at rates that were, on average, higher than in control (5.5 ± 0.7 Hz, **Figure 4A**). In agreement with previous work (Hull et al., 2013), this firing rate potentiation was preferential to Golgi cells with low baseline firing rates (FRP; <5Hz: 1.6 ± 0.2 norm, n=13; >5Hz: 1.0 ± 0.1 norm, n=4; **Figure 4B-D**). These results suggest that acetylcholine can act as an endogenous mechanism for inducing Golgi cell firing rate potentiation by activating M2 muscarinic receptors. Moreover, the diversity of plasticity across Golgi cells with different initial firing rates was in agreement with the diversity of changes in sIPSC rates in granule cells following muscarine application (**Figure 3D**).

**Figure 4.**
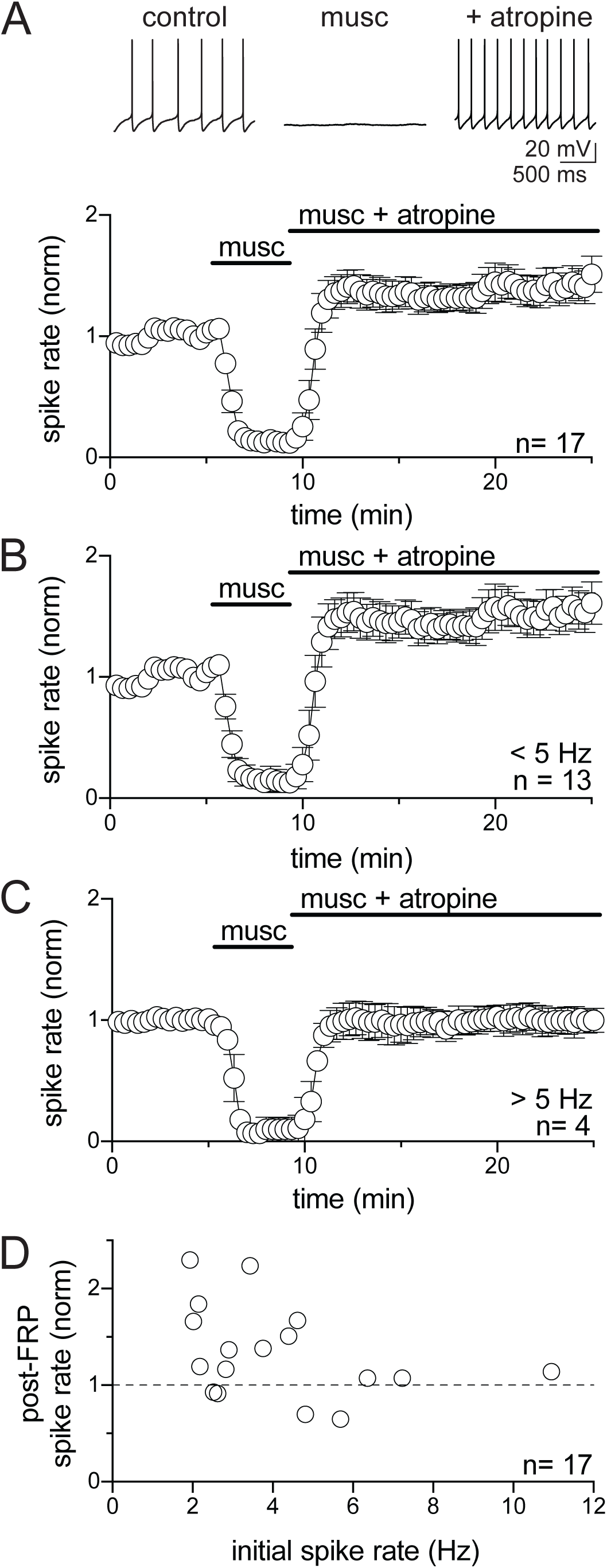
Activation of M2 receptors on Golgi cells invokes firing rate potentiation. Whole-cell, current-clamp recordings from Golgi cells before and during bath application of muscarine (musc, 10 μM). **A)** *Top*, representative current-clamp recording during muscarine application. Bath application of muscarine silences the spontaneous firing activity and this effect is reversed by the non-selective mAChR antagonist, atropine. *Bottom*, normalized spike rate across all experiments (rate at 5’: 4.2 ± 0.6 Hz; 20’: 5.5 ± 0.7 Hz; mean ± SEM; n = 17). **B)** Normalized spike rate across experiments that had a baseline firing rate below 5 Hz (rate at 5’: 3.2 ± 0.3 Hz; 20’: 4.7 ± 0.6 Hz; n = 13). **C)** Normalized spike rate across experiments that had a baseline firing rate above 5 Hz (rate at 5’: 7.7 ± 1.5 Hz; 20’: 7.9 ± 1.8 Hz; n = 4). **D)** Plot showing the initial spike rate and FRP rate for individual cells. The degree of FRP was dependent on initial spike rate.

### Cholinergic regulation of evoked granule cell inhibition

Golgi cells also provide evoked feed-forward inhibition in response to mossy fiber activation that serves to regulate excitability during cerebellar input (Duguid et al., 2015; Kanichay and Silver, 2008). To test how evoked granule cell inhibition is regulated by ongoing cholinergic modulation, we electrically stimulated the mossy fibers during bath application of muscarine. Mossy fiber stimulation reliably evoked IPSCs (control: 51.8 ± 8.4 pA, 12.8 ± 5.8% failure rate, n = 5, **Figure 5**). In the presence of muscarine, evoked IPSC amplitudes decreased significantly (98.2 ± 0.8% decrease, 0.6 ± 0.4 pA, P<0.0001, **Figure 5 B-D**), and the failure rate increased significantly (98.4 ± 0.8% failure rate, P=0.0004, **Figure 5E**). Atropine reversed these effects on the evoked IPSC amplitude (11.0 ± 15.7%, 53.0 ± 9.9 pA, **Figure 5D**) and failure rate (19.3 ± 7.1% failure rate, **Figure 5E**). Importantly, evoked IPSCs were di-synaptically evoked, as glutamatergic antagonists (NQBX 5 μM, CPP 2.5 μM) abolished all evoked IPSCs (100 ± 0% failure rate, **Figure 5E**). The muscarine-induced reduction in evoked IPSC amplitude and increase in failure rates are consistent with our finding of muscarinic receptor-mediated Golgi cell hyperpolarization. However, because these experiments rely on stimulated mossy fiber input, it is also possible that a muscarinic receptor-mediated reduction in presynaptic mossy fiber input to Golgi cells could also contribute to the observed reduction in evoked IPSCs.

**Figure 5.**
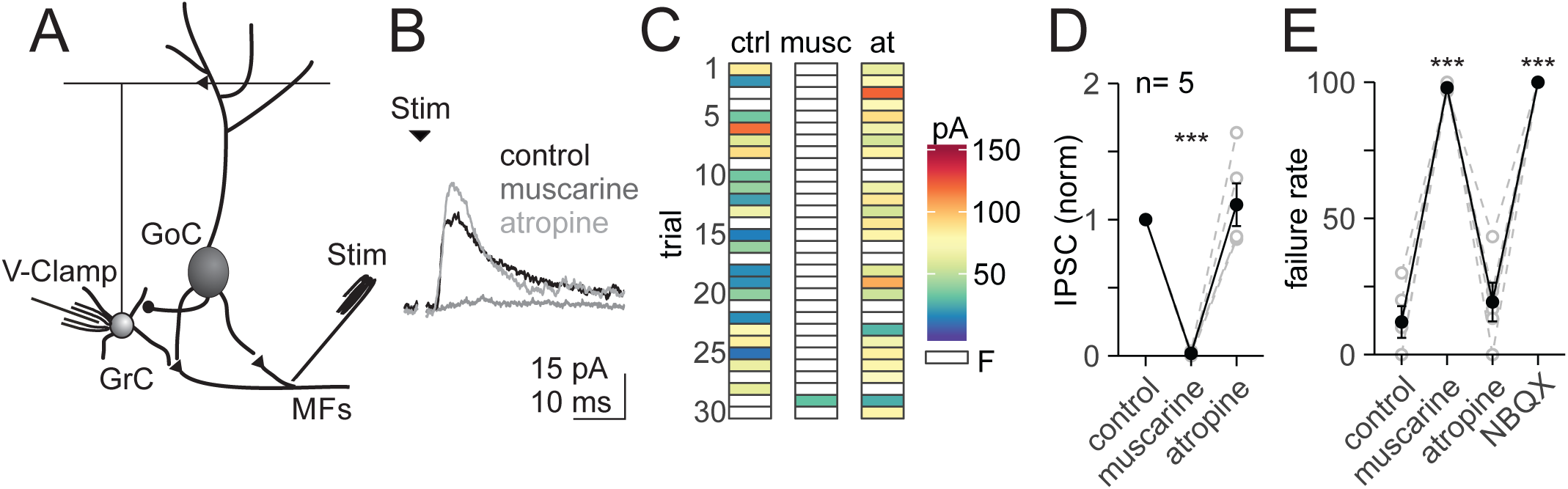
Reduction of feed-forward inhibition by muscarinic receptors. **A)** Schematic showing the intracellular recording configuration and stimulus location, ∼500–1000 µm from the cell. **B)** Representative voltage-clamp recordings of IPSCs (averaged from 30 consecutive events recorded at ∼10 mV) evoked by electrical stimulation (2 pulses at 25 Hz, 100 μs pulse width) of mossy fibers in control, muscarine (10 μM), and atropine (5 μM). **C)** Response patterns of evoked IPSCs (peak amplitude) for 30 consecutive trials in each drug condition. Failures are shown in white. **D-E)** Summary of all recorded cells. **D)** Average amplitude of the first evoked IPSC (30 consecutive trials, measurement includes successes and failures, values normalized to control) in muscarine (10 µM, 0.0 ± 0.0 norm, P<0.0001), and atropine (5 µM, 1.1 ± 0.2 norm, P = 0.7283). Individual cells are depicted in grey, black lines and error bars show mean ± SEM (ANOVA, F_1.003, 4.014_ = 44.17, P = 0.0026, n = 5). **E)** Summary of IPSC failure rate to the first stimulus in control (12.0 ± 5.83 %), muscarine (10 µM, 98.0 ± 0.8 %, P = 0.0004), and atropine (5 µM, 19.3 ± 7.1 %, P = 0.4479), and glutamate antagonist (NQBX 5 µM & CPP 2.5 µM, 100.0 ± 0.0 %, P = 0.0003; ANOVA, F_1.561, 6.244_ = 132.8, P<0.0001).

### Presynaptic Muscarinic Receptors reduce Mossy Fiber input to the granule cell layer

To test whether long-term cholinergic release could also regulate the efficacy of glutamatergic mossy fiber inputs, we next tested how bath application of muscarine regulates EPSCs onto both Golgi cells and granule cells. We first tested for the presence of muscarinic receptors at mossy fiber to Golgi cell synapses (**Figure 6**). In response to mossy fiber stimulation, muscarine significantly decreased the amplitude of evoked EPSCs onto Golgi cells (50.7 ± 11.7%, control: 127.0 ± 16.6 pA, muscarine: 61.4 ± 22.1 pA, atropine: 120.2 ± 18.7 pA, n=10, P=0.0028, **Figure 6B-D**) and increased the failure rate (control: 2.2 ± 1.0%, muscarine: 29.6 ± 8.5%, atropine: 2.2 ± 0.6%, P<0.0001, **Figure 6E**). Notably, we did not observe a significant increase in paired-pulse ratio in muscarine, as would be expected if muscarine acts presynaptically to reduce release probability (**Figure 6G**). However, mossy fibers also express GABA_B_ receptors, and muscarine silences Golgi cell spiking to reduce ongoing GABA release. Thus, muscarine should also reduce presynaptic GABA_B_ receptor activation, and this effect could serve to produce a competing increase release probability. In the presence of the GABA_B_ receptor antagonist CPG 54626 (1 μM), however, muscarine produced only a modest trend toward larger paired-pulse ratios (control PPR = 0.9 ± 0.0, muscarine PPR = 1.0 ± 0.1, p = 0.7, **Figure 6H**). We note, however, that the strong reduction in EPSC amplitude and multiple EPSC failures in muscarine add considerable variability to PPR measurements in this condition.

**Figure 6.**
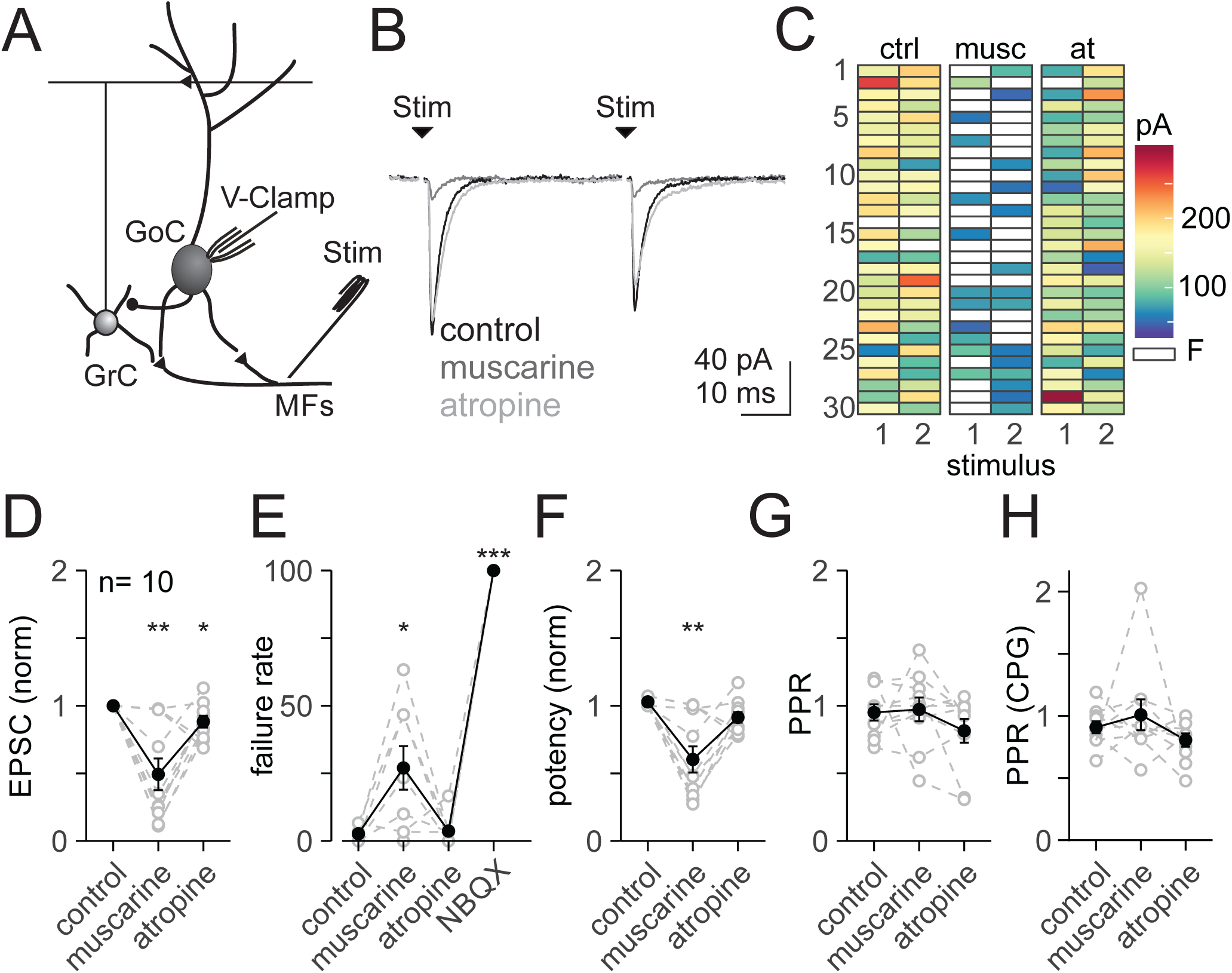
Activation of presynaptic muscarinic receptors alters glutamatergic transmission at the mossy fiber-Golgi cell synapse. **A)** Schematic showing the intracellular recording configuration and stimulus location. **B)** Representative voltage-clamp recordings of EPSCs (averaged from 30 consecutive events recorded at −60 mV) evoked by electrical stimulation (2 pulses at 25 Hz, 100 μs pulse width) of mossy fibers in control, muscarine (10 μM), and atropine (5 μM). **C)** Response patterns of evoked EPSCs (peak amplitude) for 30 consecutive trials in each drug condition. Failures are shown in white. **D-G)** Summary of all recorded cells. **D)** Average amplitude of the first evoked EPSC (30 consecutive trials, measurement includes successes and failures, values normalized to control) in muscarine (10 µM, 0.4 ± 0.1 norm, P = 0.0035), and atropine (5 µM, 0.9 ± 0.0 norm, P = 0.0439). Individual cells are depicted in grey, black lines and error bars show mean ± SEM (ANOVA, F_1.192, 10.73_=13.58, P = 0.0028, n = 10). **E)** Summary of EPSC failure rate to the first stimulus in control (2.2 ± 0.96 %), muscarine (29.6 ± 8.52 %, P = 0.0358), atropine (2.2 ± 0.6 %, P = 0.8209), and glutamate antagonist (NQBX 5 µM & CPP 2.5 µM, 100.0 ± 0.0 %, P<0.0001; ANOVA, F_1.063, 9.565_ = 120.9, P<0.0001, n = 10). **F)** Summary of EPSC potency to the first stimulus (measurement excludes event failures) in control (1.0 ± 0.0 norm), muscarine (0.6 ± 0.1 norm, P = 0.0027), and atropine (0.9 ± 0.0 norm, P = 0.0792; ANOVA, F_1.345, 12.11_=13.72, P = 0.0017). **G)** Summary of paired-pulse ratio (PPR) in control (1.0 ± 0.1), muscarine (1.0 ± 0.1 norm, P = 0.9895), and atropine (0.8 ± 0.1 norm, P = 0.5238; ANOVA, F_1.914, 17.23_=0.7338, P = 0.4889). **H)** Summary of paired-pulse ratio (PPR) in the presence of CGP (1 μM) in control (0.9 ± 0.0), muscarine (1.0 ± 0.1 norm, P = 0.7094), and atropine (0.8 ± 0.1 norm, P = 0.3975; ANOVA, F_1.492, 13.43_=1.225, P = 0.3040).

To test whether presynaptic regulation of mossy fiber input is selective to Golgi cell synapses, we next measured evoked EPSCs onto granule cells during bath application of muscarine (**Figure 7**). Again, we found a significant decrease in mossy fiber-evoked EPSCs in muscarine, along with a significant increase in failure rate (52.8 ± 9.60% EPSC decrease, control: 68.6 ± 10.7 pA, muscarine: 32.4 ± 8.8 pA, n = 10, control failure rate: 0.3 ± 0.3%, muscarine: 26.3 ± 7.5%, **Figure 7B-F**). These effects were reversed in atropine (11.8 ± 4.4%, amplitude: 64.0 ± 11.4 pA, failure rate: 1.3 ± 0.7%, **Figure 7B-F**). As with Golgi cell synapses, muscarine did not significantly increase the paired-pulse ratio (control: 0.8 ± 0.0 muscarine: 0.9 ± 0.1, atropine: 0.8 ± 0.0, P = 0.4889, **Figure 7G**). These data reveal that ongoing ACh release can reduce the efficacy of excitatory glutamatergic mossy fiber input to granule cell layer neurons by activating presynaptic muscarinic receptors.

**Figure 7.**
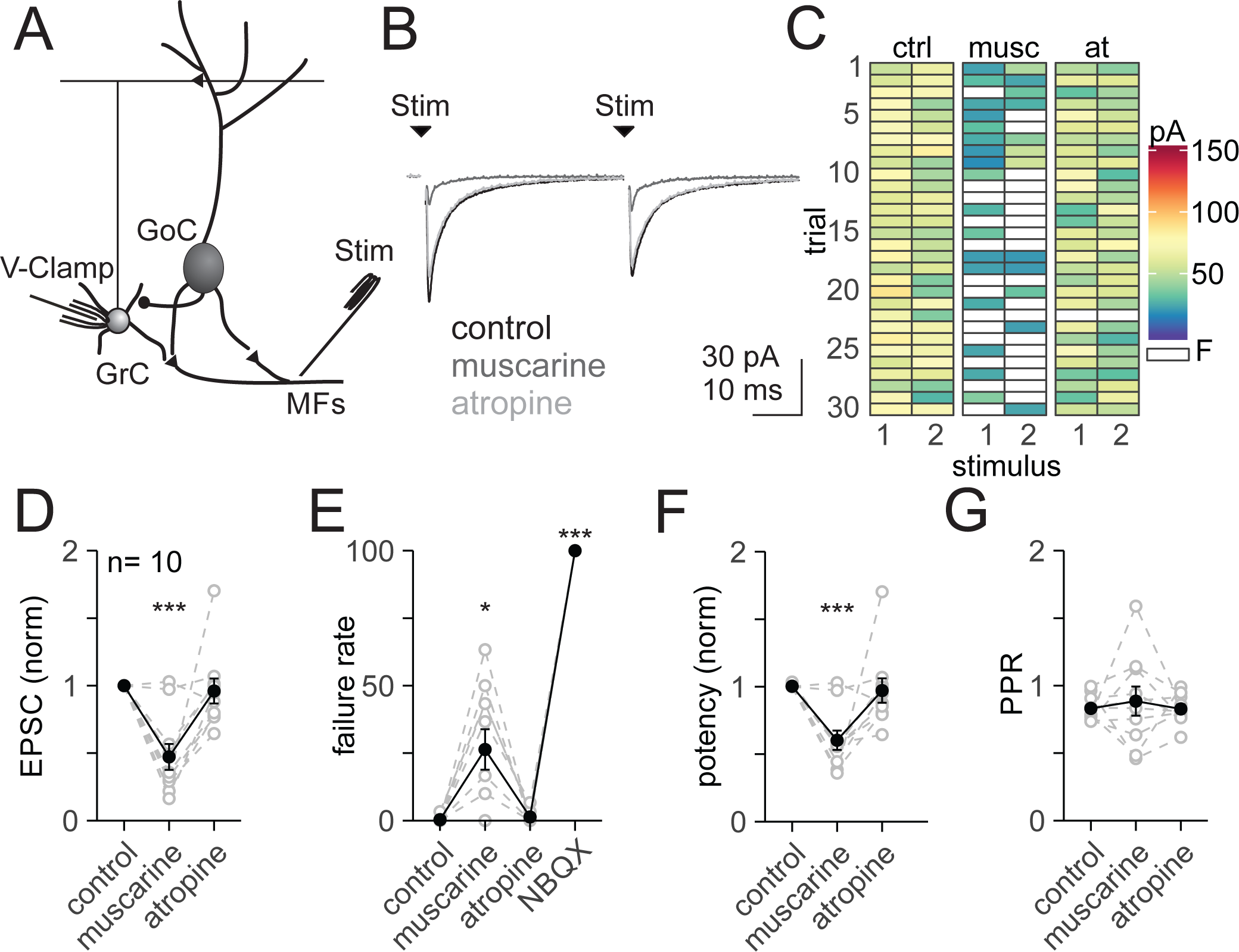
Activation of presynaptic muscarinic receptors reduces glutamatergic transmission at the mossy fiber-granule cell synapse. **A)** Schematic showing the intracellular recording configuration and stimulus location. **B)** Representative voltage-clamp recordings of EPSCs (averaged from 30 consecutive events recorded at −70 mV) evoked by electrical stimulation (2 pulses at 25 Hz, 100 μs pulse width) of mossy fibers in control, muscarine (10 μM), and atropine (5 μM). **C)** Response patterns of evoked EPSCs (peak amplitude) for 30 consecutive trials in each drug condition. Failures are shown in white. **D-G)** Summary of all recorded cells. **D)** Average amplitude of the first evoked EPSC (30 consecutive trials, measurement includes successes and failures, values normalized to control) in muscarine (10 µM, 0.5 ± 0.1 norm, P = 0.0007), and atropine (5 µM, 1.0 ± 0.1 norm, P = 0.8771). Individual cells are depicted in grey, black lines and error bars show mean ± SEM (ANOVA, F_1.65, 14.85_=14.98, P = 0.0005, n = 10). **E)** Summary of EPSC failure rate to the first stimulus in control (0.3 ± 0.3 %), muscarine (10 µM, 26.3 ± 7.5 %, P = 0.017), and atropine (5 µM, 1.3 ± 0.7 %, P = 0.4037), and glutamate antagonist (NQBX 5 µM & CPP 2.5 µM, 100.0 ± 0.00 %, P<0.0001; ANOVA, F_1.017, 9.156_ = 160.0, P<0.0001). **F)** Summary of EPSC potency to the first stimulus (measurement excludes event failures) in control, muscarine (0.6 ± 0.1 norm, P = 0.0007), and atropine (1.0 ± 0.1 norm, P = 0.9183; ANOVA, F_1.388, 12.49_=10.02, P = 0.0047). **G)** Summary of paired-pulse ratio (PPR) in control (0.8 ± 0.0), muscarine (0.9 ± 0.1 norm, P = 0.8560), and atropine (0.8 ± 0.0 norm, P = 0.9951; ANOVA, F_1.276, 11.49_=0.2322, P = 0.6969).

### Muscarinic receptor activation produces bi-directional changes in granule cell spike probability

Our results show that ACh can act through muscarinic receptors to reduce both incoming excitation and synaptic inhibition in the granule cell layer. To test how these effects combine at the circuit level to regulate granule cell activity, we performed non-invasive cell-attached recordings from granule cells while stimulating mossy fiber input (**Figure 8**). Across 8 recorded granule cells that spiked in response to mossy fiber stimulation (methods), bath application of muscarine significantly altered spike probability (**Figure 8 C-F**). Importantly, however, modulation of granule cell spiking fell into two distinct categories: In 5 of 8 granule cells, we observed an increase in granule cell spike probability and rate in response to muscarine application (muscarine: 1.9 ± 0.3 normalized rate, n = 5, p = 0.0814, **Figure 8C, D**). The remaining 3 granule cells exhibited a large decrease in spike probability and rate following muscarine application (muscarine: 0.2 ± 0.2 S.D. normalized rate, n = 3, p = 0.0612, **Figure 8E, F**). In contrast, blocking all ionotropic GABAergic inhibition (SR95531 5 μM) dramatically increased spike probabilities and rates uniformly across all recorded granule cells (muscarine: 1.2 ± 0.4 normalized, GABAzine: 6.8 ± 1.6; ANOVA F_1.076, 7.53_ = 14.95, P=0.0049, n = 8, **Figure 8C-G**). Consistent with the fast membrane time constant of granule cells (Fleming and Hull, 2019), however, neither muscarine nor SR95531 altered the timing of the first spikes in response to stimulation (mean ± S.D. stim 1: control = 3.5 ± 1.3 ms, musc = 4.3 ± 2.0 ms; stim 2: control = 2.3 ± 0.8 ms, musc = 2.5 ± 1.4 ms; stim 3: control = 2.5 ± 1.2 ms, musc = 2.8 ± 1.4 ms; n=5; **Figure 8H**). Thus, these data reveal that muscarine can alter granule cell responses by enhancing the reliability and rate of spiking in some granule cells receiving mossy fiber input, and reducing spike probability and rate in others.

**Figure 8.**
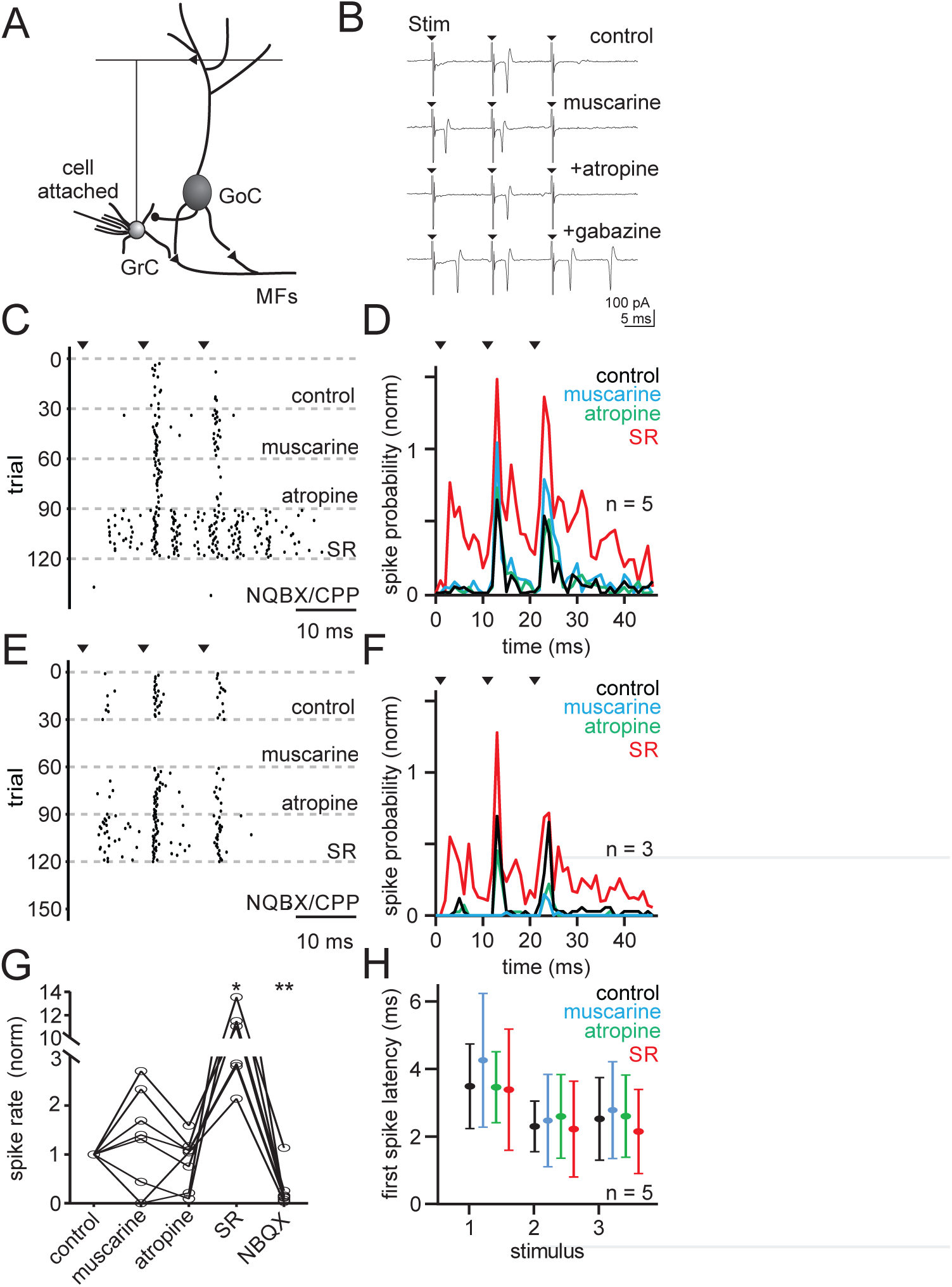
Muscarinic receptors modulate GrC spike probability. **A)** Schematic showing the cell-attached recording configuration and stimulus location. **B)** Representative cell-attached recording. Stimulus intensity was set to generate GrC spiking with approximately 20-60 % probability in response to either the 2^nd^ or 3^rd^ stimulus (3 pulses at 40Hz, repeated at 0.1 Hz). **C)** Spike raster from an example granule cell that increased spike probability in muscarine. **D)** Summary of spike probability across all granule cells that increased spiking in muscarine (n=5). Spike probability was measured by normalizing each cell to its maximum response in control. **E)** Spike raster from an example granule cell that decreased spike probability in muscarine. **F)** Summary of spike probability across all granule cells that decreased spiking in muscarine (n=3). **G)** Summary of spike rates in control, muscarine (10 μM, 1.2 ± 0.4 norm, P = 0.9087), atropine (5 μM, 0.9 ± 0.2 norm, P = 0.8923), GABA_A_ antagonist (GABAzine, 5 μM, 6.8 ± 1.6 norm, P = 0.0259), and glutamatergic antagonists (NQBX 5 µM & CPP 2.5 µM, 0.3 ± 0.1 norm, P = 0.0021; ANOVA, F_1.076, 7.53_ = 14.95, P = 0.0049, n = 8). **H)** Summary of the first spike latency in each drug condition (mean & SD). Grouped average to 1^st^ stimulus (3.8 ± 0.2 ms), 2^nd^ (2.4 ± 0.1 ms), and 3^rd^ (2.6 ± 0.1 ms; two-way ANOVA, F_3, 84_ = 1.01, P = 0.3925).

### Network model of granule cell spiking in response to muscarinic neuromodulation

How does muscarine produce this population level, bi-directional change in granule cell spiking? Because it is not possible to record sub-threshold excitation and inhibition across pharmacological conditions in the same granule cells where cell-attached recordings of spiking were performed, we generated a network model of granule cell activity to test what parameters determine whether granule cells will increase or decrease their spike probabilities in response to muscarinic neuromodulation. As muscarine reduces both excitatory and inhibitory inputs to granule cells, we expect intuitively that the net effect on granule cell firing will depend on which reduction is strongest – granule cells in which excitatory inputs are more strongly reduced than inhibitory inputs should decrease their spiking activity, while the opposite should occur in the opposite situation where inhibitory inputs are more reduced.

To test this hypothesis, we simulated a granular layer network of 4,096 granule cells and 27 Golgi cells, receiving inputs from 315 mossy fibers, with realistic connectivity parameters (methods). In this network simulation, we implemented all the effects of muscarine characterized above, in such a way as to match experimental results (methods). Upon stimulation of a random subset of 32 mossy fibers, granule cell spiking activity was qualitatively similar to the slice preparation. Of the 319 granule cells that spiked in response to mossy fiber stimulation, 85 exhibited an increase in spiking activity in the simulated presence of muscarine (peak spike probability in muscarine: 2.53 ± 0.13, normalized to control, **Figure 9 A,C**), while 65 showed decreased spiking activity (peak spike probability in muscarine: 0.33 ± 0.04, normalized to control, **Figure 9 B,C**). The remaining granule cells did not change spike probability, and were excluded from further analysis.

**Figure 9.**
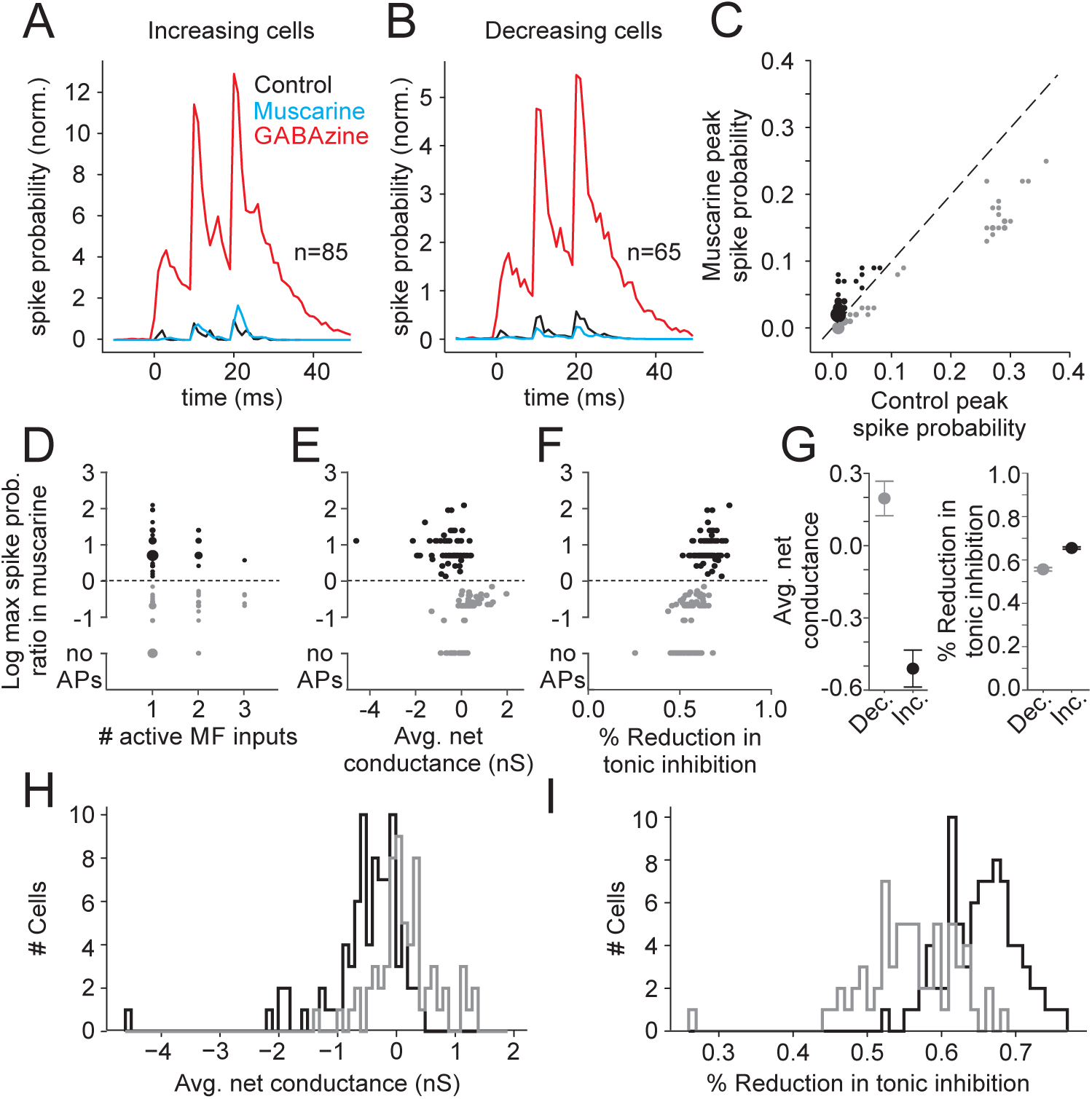
A computational model shows that the heterogeneity in granule cell responses can be explained by the variability in input connectivity and synaptic conductances. **A)** Summary of spike probability across all simulated granule cells that increased spiking in muscarine. **B)** Summary of spike probability across all simulated granule cells that decreased spiking in muscarine. **C)** Summary of peak spike probabilities for simulated granule cells from (A) and (B) in muscarine vs. control. Black denotes granule cells that increased spike probability, and gray denotes cells that decreased spike probability. **D)** Summary of granule cell peak spike probability in muscarine vs. number of active mossy fiber inputs. **E)** Summary of granule cell peak spike probability in muscarine vs. average net (excitatory – inhibitory) conductance in control conditions. **F)** Granule cell peak spike probability in muscarine vs. fractional reduction of inhibitory tonic current. **G)** Summary of the average net control conductance (left) and inhibitory tonic current (right) for simulated granule cells that increased (black) or decreased (gray) spiking in muscarine. **H)** Distributions of average net input in control for simulated granule cells that increased (black) or decreased (gray) spiking in muscarine. **I)** Distributions of fractional reduction of inhibitory tonic current in simulated granule cells that increased (black) or decreased (gray) spiking in muscarine.

We found no significant correlation between the number of active mossy fiber inputs and the change in spike probability in muscarine (Spearman’s correlation; r = 0.03 p = 0.66, **Figure 9 D**). However, cells exhibiting an increase in spike probability were dominated by inhibition in control conditions (net conductance: −0.51 ± 0.08 nS, **Figure 9 E,G**), whereas cells with decreasing spike probability received relatively less Golgi cell input (net conductance: 0.20 ± 0.07 nS, **Figure 9 E,G**). Moreover, there was a preferentially greater reduction in tonic inhibition of cells with increased spiking activity (increasing: 65.5 ± 1% reduction, decreasing: 55.7 ± 1% reduction, Kolmogorov-Smirnov D = 0.62, p < 0.0001, **Figure 9 F, G**).

We tested that our results did not depend on a specific realization of the network connectivity by simulating 30 independent realizations of the connectivity matrix, and patterns of input mossy fiber stimulation. While the relative numbers of cells exhibiting increases and decreases in spike probability were variable across different instantiations of the model, the underlying input profiles distinguishing the two cell categories were comparable (increasing cells had a net inhibitory conductance in control: −0.40 ± 0.03 nS, decreasing cells had a net excitatory conductance in control: 0.26 ± 0.03 nS, not shown). Similarly, we saw consistently larger decreases in the inhibitory tonic current between for cells that increased their spike probability in muscarine (increasing: 64.7 ± 0.2% reduction, decreasing: 57.6 ± 0.2% reduction, not shown). Together, these results indicate that muscarinic neuromodulation preferentially enhances the spike rates of granule cells with larger synaptic inhibition in control conditions.

## Discussion

Here we have shown that acetylcholine activates muscarinic receptors to modulate two of the three main nodes within the granule cell layer of the cerebellar cortex. Specifically, both Golgi cells and mossy fibers express muscarinic receptors that serve to hyperpolarize Golgi cells and decrease mossy fiber release probability respectively. By simulating ongoing release of acetylcholine using bath application of muscarine, we revealed a suppression of Golgi cell spiking that is mediated by M2-type muscarinic receptors. Consistent with this finding, optogenetic stimulation of ChAT+ cerebellar projections produced a long-lasting, net hyperpolarizing current in Golgi cells that greatly outlasted the duration of stimulation. At the circuit level, suppression of Golgi spiking led to a large decrease in synaptic inhibition onto granule cells. In parallel, muscarine also reduced excitatory transmission from mossy fibers by acting on presynaptic muscarinic receptors.

Together, by acting on both Golgi cells and mossy fibers, we find that ACh can alter the coordination of excitation and inhibition onto granule cells. Surprisingly, however, these changes in excitation and inhibition produce diverse changes in granule cell spiking at the population level. Specifically, spike probability in some granule cells can be enhanced, while others are largely suppressed. By generating a network model of the granule cell layer, we provide evidence that the direction of spike rate modulation is established by the initial balance of excitation and inhibition onto each cell, as well as the degree to which neuromodulation reduces tonic inhibition. Overall, cells with higher levels of inhibition increased their spike rates in response to neuromodulation, while cells with less inhibition decrease their spike rates. These changes are in stark contrast to the responses observed after removing all inhibition, which dramatically increases spiking of the entire population. Thus, these data suggest that by simultaneously reducing excitation and inhibition, cholinergic neuromodulation can provide a mechanism for enhancing the responses of subsets of granule cells without expanding the overall population response.

Under what conditions might ACh act to modulate the granule cell layer in vivo? Previous work suggests that ChAT+ cerebellar projecting neurons originate from diverse sources (Jaarsma et al., 1997). Consistent with our findings, these projections can take the form of mossy fiber-like inputs, or thinner beaded fibers (Jaarsma et al., 1997). Previous studies have shown that ChAT+ mossy fiber terminals and beaded fibers are expressed throughout the cerebellar cortex, with lobules IX/X having the densest mossy fiber innervation and lobule IX having the densest beaded fiber innervation (Barmack et al., 1992a; Jaarsma et al., 1996; Ojima et al., 1989). We also observed the densest innervation of ChAT+ mossy fiber-like terminals in these lobules, though ChAT+ terminals were present in all lobules, and we observed optogenetically driven currents in Golgi cells throughout the vermis. Anatomical studies have suggested that the ChAT+ MFs located in lobules IX/X and the flocculus originate from medial vestibular nucleus (MVN) and nucleus prepositus hypoglossi (NPH) (Barmack et al., 1992b), whereas ChAT+ beaded fibers may originate from the pedunculopontine tegmental nuclei (PPTg) (Newman and Ginsberg, 1992; Ruggiero et al., 1997; Woolf and Butcher, 1989). Notably, for any of these sources it remains possible that other neurotransmitters are co-released with ACh as has been shown in other brain regions (Granger et al., 2017). However, based on our goal of measuring the longer timescale effects of ACh, we did not investigate the possibility of other fast transmitters that may also play a role during the initial phases of transient ACh release.

Such diversity of sources of ACh suggests multiple roles for cholinergic signaling. Inputs from the medial vestibular nucleus may be important for modulation of vestibular guided reflexes. In support of a role for ACh in modulating vestibular guided behaviors, previous work has shown that cerebellar injection of cholinergic agonists can enhance both OKR and VOR reflexes (Prestori et al., 2013; Tan and Collewijn, 1991, 1992). Such findings are in line with an enhancement of responses in granule cells, and particularly those receiving vestibular input. Thus, the increased spike probabilities observed here in a subpopulation of granule cells may explain these previously observed behavioral effects of ACh.

The presence of cholinergic inputs from the PPTg also suggest a role for ACh in contextual modulation of the granule cell layer, for instance by arousal or locomotion. Indeed, recent work has demonstrated a locomotion-dependent enhancement of learning in a cerebellar dependent form of delay eyeblink conditioning (EBC) (Albergaria et al., 2018). Specifically, the acquisition rate of delay EBC was correlated with the animals running speed. In this study, the circuit mechanism underlying this enhancement was linked to an increase in granule cell layer excitability (Albergaria et al., 2018). In this model, behaviorally relevant sensory information would more easily drive granule cells past spike threshold during locomotion, and thus transmit more effectively to downstream Purkinje cells to promote synaptic plasticity and learning. While speculative, our results suggest the possibility that ACh could provide an endogenous mechanism to disinhibit populations of granule cells and enable such locomotion-dependent effects on learning. In particular, because the PPTg is active during locomotion (Lee et al., 2014), it could provide the type of tonic acetylcholine release that would enhance granule cell responses and promote associative learning. At present, however, it is unknown whether ACh is released into the cerebellar cortex during locomotion.

Finally, our results suggest that the key determinant of how ACh modulates granule cell spike probability is the initial balance of excitation and inhibition onto each cell. Indeed, previous work has demonstrated that inhibition is not homogenous across granule cells (Crowley et al., 2009). Such diversity may arise from a combination of factors, including differences in the number of Golgi cell inputs per glomerulus (Jakab and Hamori, 1988), the number of granule cell dendrites per cell (Palay and Chan-Palay, 1974), a diversity of Golgi cell types (Geurts et al., 2001; Neki et al., 1996; Simat et al., 2007), or differences in GABA receptor expression at granule cell synapses (Wall, 2002). Likewise, the amplitude of unitary excitatory mossy fiber conductances onto granule cells has been shown to vary over a wide range, and depend on the source of the mossy fiber input (Chabrol et al., 2015). Moreover, because granule cells can integrate multiple mossy fiber inputs from diverse sources (Huang et al., 2013), it is possible for individual cells to exhibit considerable variability in both the amount of excitation and inhibition they receive across the population.

Notably, we find ACh selectively increases the excitability of the granule cells that are most strongly inhibited. Previous work has suggested that that granule cell inhibition can be stimulus-specific (Precht and Llinas, 1969), and recent work has indicated that such stimulus specific inhibition may play a key role in cerebellar learning (Kalmbach et al., 2011). If different Golgi cells can indeed be recruited by some stimuli and not others, cholinergic neuromodulation would provide an ideal mechanism to enhance the responses of granule cells that would otherwise be suppressed by stimulus-specific inhibition. Hence, such a mechanism could serve to enhance learning for specific mossy fiber inputs, perhaps based on behavioral context or stimulus salience. Thus, ACh neuromodulation would provide an appealing means to regulate learning or modify the gain of cerebellar transformations. While future *in vivo* experiments will be necessary to evaluate this possibility, our results reveal both the cell autonomous and circuit level actions by which ongoing ACh release can act through muscarinic receptors to modulate granule cell excitability. Such data thus provide the foundation for interpreting the *in vivo* effects of cholinergic neuromodulation at the input stage of cerebellar processing.

## Acknowledgements

This work was supported by grants from the NIH NINDS (5R01NS096289-02), the NSF (DGF1106401) the Sloan Foundation, and the Whitehall Foundation. We thank Dr. Lindsey Glickfeld and members of the Hull and Glickfeld labs for input and technical assistance throughout the project.

